# Simulation of evoked responses to transcranial magnetic stimulation using a multiscale cortical circuit model

**DOI:** 10.1101/2025.09.02.673790

**Authors:** Zhihe Zhao, Aman S. Aberra, Alexander Opitz

**Affiliations:** Department of Biomedical Engineering, University of Minnesota, MN; Department of Biological Sciences, Dartmouth College, NH

**Keywords:** Neuronal network models, Motor cortex, Simulation, Transcranial magnetic stimulation

## Abstract

**Background:** Transcranial magnetic stimulation (TMS) is a widely used non-invasive brain stimulation technique, but the neural circuits activated by TMS remain poorly understood. Previous modeling approaches have been limited to either simplified point-neuron networks or isolated single-cell models that lack synaptic connectivity.

**Objective:** To develop and validate a multiscale cortical circuit model that integrates morphologically-realistic neurons with accurate TMS-induced electric field distributions and to investigate mechanisms underlying cortical responses to stimulation.

**Methods:** We constructed a network model of a cortical column comprising 10,000 biophysically realistic neurons (excitatory pyramidal cells and inhibitory interneurons) across layers 2/3, 5, and 6 with over 10 million synaptic connections. The model incorporated thalamic and non-specific corticocortical inputs to generate physiological firing rates. TMS-induced electric fields were calculated using finite element modeling and coupled to individual neurons through the extracellular mechanism. We validated model predictions against experimental recordings of TMS-evoked local field potentials (LFPs) and multiunit activity.

**Results:** The model reproduced key experimental observations including the dose-dependent N50 LFP component and multiphasic multi-unit responses, consisting of an (excitatory) increase followed by a (inhibitory) decrease in firing rates. The early excitatory response exhibited dual-peak dynamics reflecting distinct contributions from directly and indirectly activated neuronal populations, and the subsequent inhibitory phase reflected activation of feedback GABAergic circuits through both GABA_A_ and GABA_B_ conductances. Spatial analysis across 30 cortical columns distributed across the precentral gyrus revealed orientation-dependent evoked responses.

**Conclusion:** This validated multiscale model provides mechanistic insights into TMS-evoked cortical dynamics, demonstrating how direct neuronal activation cascades through synaptic networks to generate characteristic population responses. The framework establishes a computational platform for optimizing stimulation protocols in research and clinical applications.

**Graphical Abstract:** 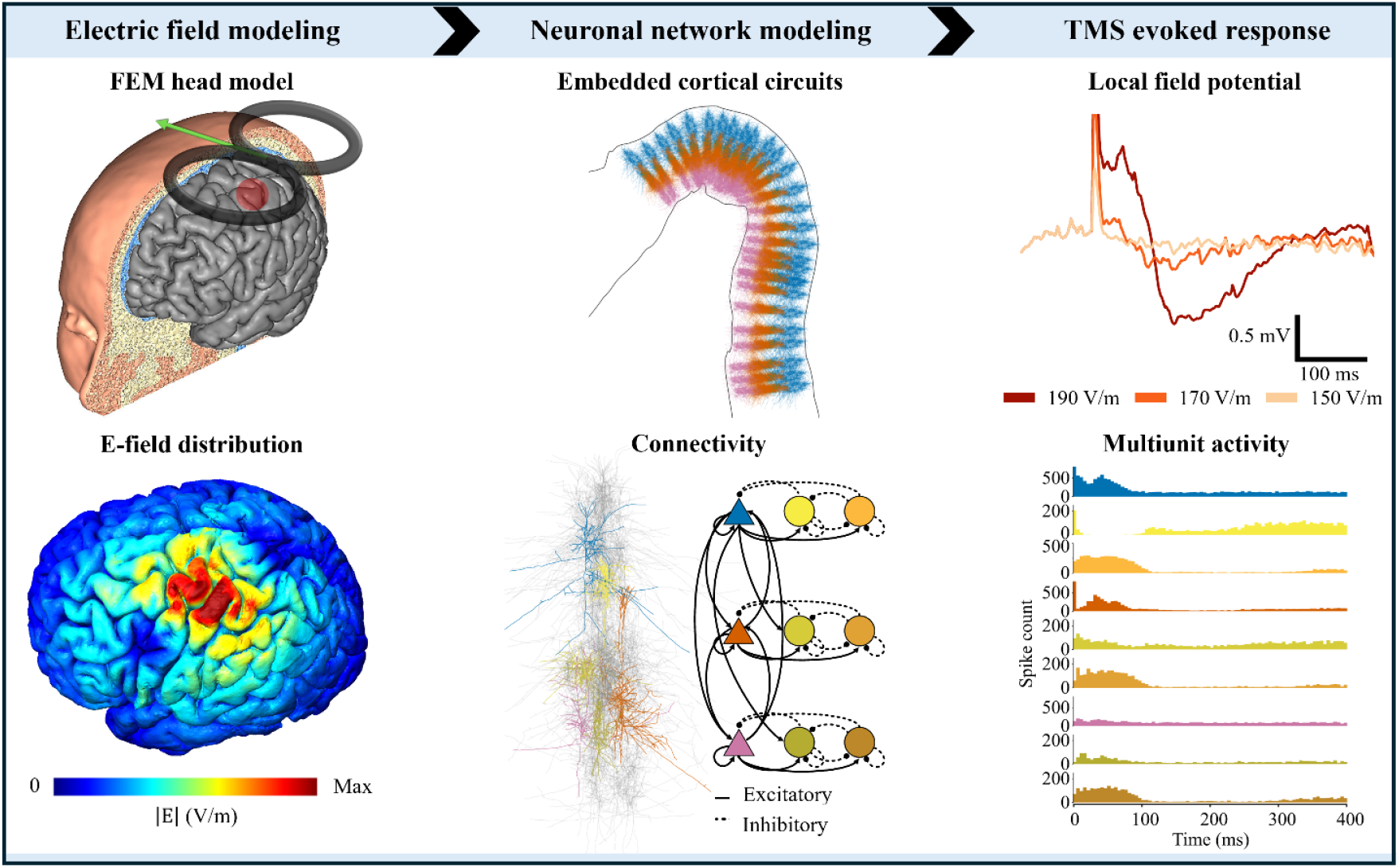

## Introduction

Transcranial magnetic stimulation (TMS) is a non-invasive brain stimulation technique widely used in research and clinical applications. In TMS, a time-varying magnetic field is applied to the head which induces an electric field (E-field) in underlying tissue and can directly activate neurons in the brain. TMS is FDA-approved for treatment-resistant depression, obsessive-compulsive disorder, and nicotine addiction and is being explored for other psychiatric disorders [1,2]. Despite its increasing use in research and clinical applications, the neural circuits activated during TMS remain poorly understood. Computational neural modeling provides a powerful tool for investigating the mechanisms of TMS and optimizing stimulation parameters for stronger and targeted physiological effects. The development of TMS modeling approaches has evolved through distinct stages, each addressing different scales of neural organization.

Initially, finite element method (FEM) head models enabled accurate calculation of TMS-induced E-field distributions in realistic head geometries, providing the foundation for understanding the spatial patterns of E-field exposure across cortical regions. Building upon this E-field modeling framework, Aberra and colleagues [3–5] advanced multi-scale models by incorporating morphologically-realistic cortical neurons within a head model to simulate cellular responses to TMS-induced E-fields. This work demonstrated the importance of considering both detailed neuronal morphologies and accurate E-field distributions for predicting neuronal activation patterns and established critical relationships between E-field strength, orientation, and neuronal activation thresholds. However, this single-neuron modeling approach was limited to examining responses of isolated neurons without considering their synaptic interactions within cortical circuits, which are essential for understanding the network-level effects of TMS.

Earlier simulation studies using network models with simplified point neurons successfully reproduced certain characteristics of TMS-evoked corticospinal activity [6,7]. However, these models do not explicitly represent the interaction between the induced E-field and neurons. Instead, they rely on assumptions about the number, location, and connectivity of directly activated neurons. This disconnect between detailed single-neuron multi-scale modeling and network-level dynamics represents a critical gap in our understanding of how TMS affects cortical circuits. The next essential step in this modeling pipeline requires building multi-scale models that integrate morphologically-realistic neurons, accurate E-field coupling, and their synaptic connectivity within realistic cortical networks.

Here, we develop a novel multi-scale computational cortical circuit model that integrates morphologically-realistic neurons with accurate TMS-induced E-field distributions. Our approach bridges single-cell modeling and network-level dynamics by incorporating synaptic connectivity within a cortical column. We validate the model against experimental TMS-evoked potentials (TEP) and use it to dissect the mechanisms underlying cortical responses, revealing distinct contributions of direct versus indirect neuronal activation. By embedding multiple cortical columns within a realistic head geometry, we investigate spatial activation patterns and coil orientation effects. This integrated modeling framework provides mechanistic insights into TMS circuit dynamics and establishes a validated platform for optimizing stimulation protocols.

## Methods

### Cortical neuron models

We implemented a computational circuit model comprising a population of biophysically-based, morphologically-realistic multi-compartment cortical neurons with Hodgkin-Huxley-based ion channel dynamics, adapted from the Blue Brain network [8] and implemented in NEURON-Python 8.0 [[9]. These cells were originally reconstructed from acute slices of rat somatosensory cortex. Our circuit model incorporated three different pyramidal cell types as modified by Aberra et al. : Layer 2/3 pyramidal cells (L2/3 PC), Layer 5 thick tufted pyramidal cells (L5 TTPC), and Layer 6 tufted pyramidal cells (L6 TPC) [10]. Additionally, we modeled two principal classes of GABAergic interneurons: parvalbumin-expressing (PV) fast-spiking neurons and somatostatin-expressing (SST) low-threshold spiking neurons. For the PV interneurons, we used large basket cells (LBC) from Layers 2/3, 5, and 6 of the Blue Brain cell library [8], selected for their potentially low TMS thresholds [10]. Because the morphologically-realistic neuron models provided were from rats, they were modified and adapted to the biophysical and anatomical properties of adult human-like cortical neurons, which included geometric scaling of diameters and dendritic lengths and incorporation of myelinated axonal arbors. We followed the scaling procedure and parameters given in [11,12] to modify the LBC models. In contrast, for SST interneurons, we employed simplified neuron models [13], as they are less likely to be myelinated compared to PV interneurons [14], making them less susceptible to direct TMS activation.

### Cortical column composition: Neuron locations, densities and ratios

We modeled a cylindric volume of a human cortical column in primary motor cortex (M1) with a diameter of 400 µm and a cortical thickness of 3000 µm. The column diameter chosen falls into the reasonable range from 300 μm to 600 μm of measurement and other existing cortical column models [8,15–18]. The overall M1 cortical thickness was derived from the BigBrain 3D Human Brain Model [19]. Each layer’s approximate relative thickness in the column were estimated as follows: L1-10%, L2/3-36.6%, L4-5.8%, L5-22.9%, L6-24.7%. Since M1 has been considered as an agranular cortex lacking layer 4[20], we excluded neurons in Layer 4 in the network. Although direct measurements of human M1 density were less frequently reported in the literature than for nonhuman primates (NHP), human M1 has a similar magnitude of neuronal density to other primates[21]. The neuronal density in motor cortex of NHP is typically in the range of about 20,000 to 30,000 neurons/mm^3^ [22–24]. In total, we have 10,000 neurons with uniform cell density of 26,525 neuron/mm^3^, which lies well within this range. The proportion of the excitatory neurons to inhibitory neurons across all layers were estimated as 4:1. The ratio of PV to SST neurons per layer was estimated as 2:1 which is consistent with a recent mouse M1 microcircuit model [18].

### Local connectivity

Local connectivity parameters were adapted from previous work by [18,25,26]. Connection probabilities between neuronal populations *Ca* were taken from these studies, and the total number of synaptic connections *K* was calculated using Peters’ rule [25–27],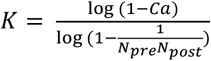, where *N*_*pre*_ and *N*_*post*_ are the number of neurons in the pre and postsynaptic population. Pre-post pairs were randomly selected with uniform probability, allowing for multiple synaptic contacts between any pair of connected neurons. The cortical column model contains over 10 million synaptic connections made within and between layers. Neurons in layers 2/3 sent strong descending projections to lower layers, particularly layer 5 [28]. All synapses were modeled as dual-exponential conductances. Excitatory synapses consisted of colocalized AMPA (rise, decay time constants τ: 0.2, 1.7 ms) and NMDA (rise, decay τ: 2, 26 ms) receptors, both with reversal potentials of 0 mV [29]. The ratio of NMDA to AMPA receptors was 1:1 [30], i.e., each accounting for 50% of the connection weight. Inhibitory synapses from SST to excitatory neurons consisted of a GABAA receptor (rise, decay τ: 0.3, 2.5 ms) and GABAB (45.2 ms, 175.16 ms) receptor, in a 0.9/0.1 ratio[18]; synapses from PV to other neurons consisted of a GABAA receptor [18]. The reversal potential was −70 mV for GABAA and −90 mV for GABAB receptors. Connection delays were 1 ms plus an additional delay based on the distance between presynaptic and postsynaptic neuronal somata, assuming a conduction velocity of 0.5 m/s. PV interneurons made synapses near the soma of excitatory neurons, while SST interneurons targeted the apical dendrites of excitatory neurons, in accordance with the literature [18,31–33]. When these interneurons (both PV and SST) themselves received inputs, those inputs arrived at their dendrites. All synapses in the model were distributed randomly with a uniform distribution.

### Long-range input connectivity

We adapted the implementation of long-range input from the mouse M1 circuit by Dura-Bernal et al. [18]. The cortical column received long-range input modeled as Poisson processes from motor thalamus and non-specific corticocortical inputs to achieve physiologically-realistic baseline firing rates (FR). The motor thalamus consisted of a population of 1000 [34,35] spike-generators (NEURON NetStims) that generated independent, Poisson-random spike trains with fixed average spike rates uniformly distributed between 0 and 5 Hz [36,37], [18]. The proportion of the excitatory to inhibitory population in the motor thalamus was estimated to be 4:1. The thalamic inputs target neurons in layers 2/3 through 5 [38]. Non-specific inputs were modelled as Poisson spike trains with a rate of 8 Hz, which target all layers (Fig. 1C). Synapses from long-range input uniformly distributed across dendrites of the postsynaptic cells.

**Figure 1.**
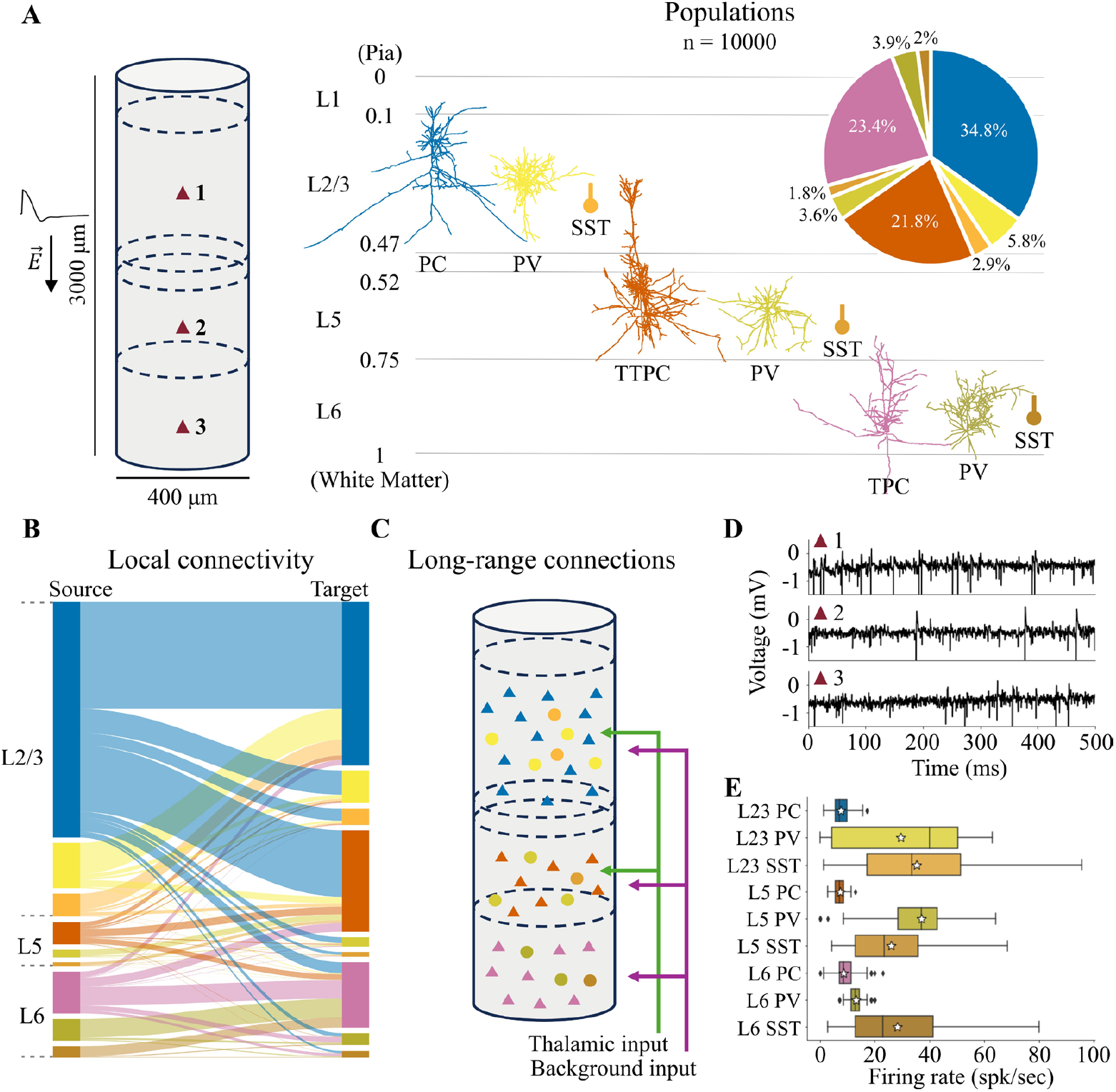
Cortical column model: dimension, population, classes, morphologies, connectivity and spontaneous activity. **(A)** Dimensions of simulated cylindrical volume (left) with three simulated recording sites and cell types and populations (right). Left shows L1–L6 boundaries with normalized cortical depth from 0 = pia to 1 = white matter. The cortical circuit consists of interconnected multi-compartmental biophysically realistic pyramidal cells (PC), realistic Parvalbumin-expressing large basket cells (PV) and reduced Somatostatin (SST) neurons. Proportion of cells in each population is shown in the pie chart. Uniform electric field is applied along the vertical axis of the column with a monophasic TMS waveform. **(B)** Local connections between populations across and within layers. Line thickness is indicative of the relative number of connections from the source to the target population. **(C)** Long-range connections from the motor thalamus (thalamic input) and other cortical and subcortical region (background input). **(D)** Simulated raw LFP at each recording site in L23, L5 and L6 (Top) and FR (Bottom, boxplots) for different cell types and layers during spontaneous firing. Boxplots show the minimum, lower quartile, median and mean (white star), upper quartile, and maximum values. **(E)** FR for different cell types in the model, shown in boxplots. The white star shows the mean.

### Electric field modeling

We calculated TMS-induced E-fields using the volume conductor head model (See Ref. [39] for modeling details) in SimNIBS v3.2, a freely available software suite that incorporates MRI image segmentation, computational mesh development, and finite element method-based E-field simulation [40]. To obtain finer E-field distribution at the M1 hand knob, we remeshed the M1 model with higher resolution, resulting in a mesh with approximately 1.6 million nodes and 9 million tetrahedral elements. E-field distributions were calculated using SimNIBS models of the Magstim 70 mm figure-of-8 coil positioned over the primary motor cortex. The coil was oriented to induce currents perpendicular to the central sulcus at 45° relative to the midline, with monophasic stimulation applied in both posterior–anterior (P–A) and anterior–posterior (A–P) directions.

### Embedding cortical column

A region of interest (ROI) containing the M1 hand knob on the precentral gyrus and part of the postcentral gyrus was selected for cortical column placement. Thirty surface mesh elements were selected on the gray matter surface for embedding and orienting the columns. In each of the 30 elements, a cortical column model was embedded by aligning the column’s vertical axis to the normal vector at the center of the element. The depth of each cortical column was determined by measuring the distance between the gray and white matter surfaces along the element’s normal vector. Following established cortical column structure[41], model neurons were oriented with their somato-dendritic axes aligned to the element normal vector and randomly rotated around these axes in the azimuthal plane to reflect biological variability. Mesh generation and extraction of E-field vectors from the SimNIBS output and neuron placement, simulations (in NEURON), and subsequent analysis were conducted in Python v3.8. Visualization was conducted using both Python and MATLAB (R2023a Mathworks, Inc., Natick, MA, USA).

### Coupling of electric fields into neuron models

Using the quasi-static approximation[42], the TMS-induced E-field was decomposed into separate spatial and temporal components. As a result, the spatial pattern of the E-field remains constant and the amplitudes at each location can be scaled over time by the E-field pulse waveform. The E-field amplitudes at each location were calculated in SimNIBS and linearly interpolated at the coordinates of the neuron model compartments. Quasipotentials [5,43] were computed for each neuron by numerically integrating the E-field along the path of each neural process. These values were then applied as extracellular scalar potentials using the extracellular mechanism, following the previously established approach for coupling TMS-induced E-fields to cable models in NEURON [11,43].

### Local field potentials (LFP) calculation

The LFP signal was obtained by summing the extracellular potential induced by each segment of each neuron at the electrode location. Extracellular potentials were calculated using the line source approximation (LSA) [44–46] and assuming an Ohmic medium with conductivity *σ*= 0.275 S/m, matching the gray matter conductivity used in the SimNIBS head model. LSA implementation has been shown to better approximate extracellular signals than point source approximation [45]. The LFP of the network was sampled at 5000 Hz.

### Neural data analysis

To analyze the LFPs, we preprocessed the signals by removing and interpolating the first 25ms post-stimulation, as this time window corresponds to the period in experimental recordings where analysis is typically not feasible due to stimulation artifacts. We then applied band-pass filtering using a 2nd-order Butterworth filter with cutoff frequencies of 1 Hz and 50 Hz. The N50 component was then extracted as the peak amplitude from the 40–70 ms window post-TMS onset [47]. For multiunit activity (MUA) analysis, normalized population FR under different stimulation intensities were calculated by subtracting the baseline average FR (no stimulation condition) from the instantaneous FR of each time bin, then averaged across neurons. The instantaneous firing rate was calculated by counting the number of spikes within time bins of 2ms, divided by the bin size and the number of neurons in the population. Significance thresholds (p<0.05) for excitatory and inhibitory events were determined using the 2.5th and 97.5th percentiles of the normalized FR distribution during the 100 ms pre-TMS period. Normalized FR traces were smoothed using a Gaussian kernel (σ = 2 ms) [48].

### Parameter exploration/optimization

We ran batch simulations to explore and optimize parameters. We specified the range of parameters and parameter values to explore, and the tool automatically submitted simulation “jobs” on a high-performance computing cluster (HPC) using the SLURM job scheduler. We primarily employed the SLURM scheduler at the Minnesota Supercomputing Institute (MSI). Simulating one second of activity in the full cortical column model using NEURON with a time step of 25 µs required approximately 30 wall-clock hours using parallelization across 50 CPU cores, resulting in a total computational cost of ∼1,500 core-hours.

Computation of each neuron was distributed across cores in a round-robin fashion. Since we performed large-scale parameter sweeps and model optimization, the total computational demand scaled significantly, requiring substantial HPC resources. Building the model and obtaining the results shown required over 2 million computational core-hours.

## Results

We developed a cortical column model comprising 10,000 interconnected neurons with either morphologically-realistic or reduced representations. To simulate spontaneous activity, we applied both thalamic and non-specific corticocortical synaptic inputs to all neurons in the model. We then computed extracellular signals from three recording locations positioned in L2/3, L5, and L6 (Fig. 1D). The population mean and median firing rates (FR) of pyramidal cells (PCs) and inhibitory interneurons ranged between 7-10 Hz and below 40 Hz, respectively. Specifically, PCs exhibited the following FR (median, interquartile range): L2/3 PCs (7.14 Hz, 4.29 Hz), L5 PCs (7.14 Hz, 2.86 Hz), and L6 PCs (8.57 Hz, 4.29 Hz) (Fig. 1E).

Applying a single TMS monophasic pulse with a 190 V/m uniform E-field vertically downward to the cortical column model revealed distinct cell-type and layer specific temporal responses (Fig. 2). Among the excitatory cells, L2/3 PC and L5 PC exhibited strong, immediate activation with synchronized firing after stimulation onset, appearing as vertical bands on the raster plot, consistent with their low TMS thresholds previously demonstrated in [10]. During the early phase (10–100ms), both L2/3 and L5 PCs showed sustained elevated firing that gradually dispersed, transitioning from synchronized to more scattered spike times. Among interneurons, PV cells in deeper layers (L5 and L6) showed elevated firing, while L2/3 PV cells exhibited suppressed activity with notably reduced FR. SST interneurons in all layers demonstrated strong elevated activity throughout this early phase. In the late phase (>100ms), most populations showed firing activity that returned to near-baseline levels. However, L5 PC and SST interneurons exhibited a distinct suppression of activity from 100 to 300ms, dropping below their own baseline FR and remaining lower than other cell types, before eventually recovering. This suppression was not observed in L2/3 PC, L6 PC, or PV interneurons, which maintained relatively stable low-level activity throughout the late phase.

**Figure 2.**
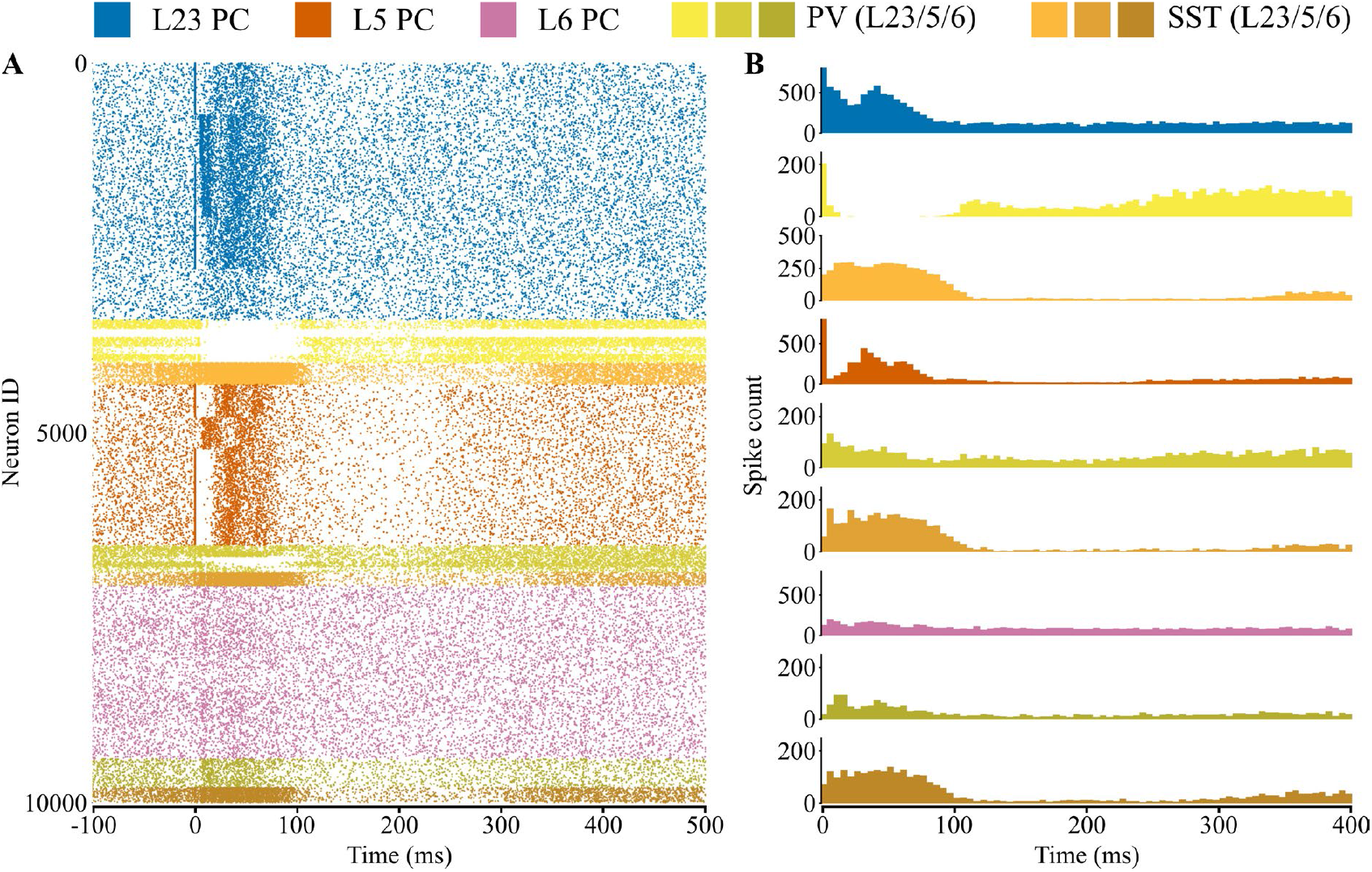
Cell-type- and layer-specific response to single monophasic TMS pulse with uniform E-field (190 V/m). **(A)** Raster plot of evoked activity (100ms before and 500ms after the TMS onset at 0ms). Cells are grouped by population. **(B)** Spike count histogram for different cell types (5ms time bins).

To investigate how direct and indirect neuronal activation contributes to the TMS evoked response, we categorized PCs into two groups based on spike latency: “direct” neurons that spiked within 1.5 ms of the TMS pulse (reflecting direct E-field activation), and “indirect” neurons that spiked after 1.5 ms, presumably due to synaptic transmission from directly activated neurons. At 150 V/m, TMS directly activated 25.5% of L5 PCs while activating minimal numbers of L2/3 PCs and L6 PCs (Table S1). At 190 V/m stimulation intensity, TMS directly activated PCs in a layer-specific pattern: L2/3 PC showed 27.5% direct and 72.5% indirect activation; L5 PC exhibited the highest proportion of direct activation at 63.85% direct and 36.15% indirect, while L6 PC had minimal direct activation with only 2.05% direct and 97.95% indirect activation (Supplementary Fig. 1). During the early phase (10–100 ms), “direct” and “indirect” L2/3 and L5 PCs exhibited alternating increases and decreases in spiking activity. Moreover, the peak FR of “indirect” L5 PC neurons occurred later than those of L2/3 PC indirect neurons. In contrast, the direct population of L6 PCs exhibited more variable FR throughout the simulation, while the indirect population FR remained relatively stable with less fluctuation.

We compared the model and experimental TMS-induced evoked responses to validate the model output. First, we compared the evoked LFP computed from L5 of the column. Specifically, the evoked LFPs showed a maximum deflection approximately 50ms after TMS onset during the early phase. This N50 component demonstrated a distinct increase in magnitude with increasing stimulation intensity, showing a robust response at higher stimulation intensity (190 V/m) (Figure 3A). This dose-dependent behavior was also observed in the later phase, where the LFPs showed a prolonged deflection with opposite polarity that increased in magnitude before returning to baseline. The dose-dependent evoked potentials closely match TEPs recorded in NHP (rhesus macaque) [47] (Figure 3A, bottom). To assess the robustness of the dose-dependent response, we ran simulations with two additional random seed sets for the background activity from long-range inputs, effectively creating three independent trials. The dose-dependent pattern remained consistent across all trials (Supplementary Fig. 2), confirming that the observed relationship was independent of the specific random seed used. Second, we compared the multi-unit activity post-stimulation from neurons in L5 (Model: L5 PC, Experiment: Layer V of rodent caudal forelimb area (CFA) [48]). Figure 3B illustrates the evoked normalized population FR (instantaneous population FR minus baseline average FR). Significance thresholds were determined based on the 2.5th and 97.5th percentiles of the normalized FR during baseline, following the approach in [48] for better comparison. Both the model and the experimental recording showed a similar multiphasic pattern of neuronal activity. In the model, after the initial synchronized firing due to direct activation, the L5 PCs exhibited elevated firing rates with a first peak around 30 ms and a second peak around 60 ms, followed by a prolonged phase of inhibition. The first and second peaks of L5 PC normalized FR corresponded to peaks in the “direct” and “indirect” neuron FR, respectively (Supplementary Fig. 1), indicating that the two excitatory peaks during the early phase reflect distinct contributions from ‘direct’ and ‘indirect’ neuronal populations through transsynaptic excitation. The amplitude and duration of both excitatory and inhibitory phases in L5 PC FR scaled with stimulation intensity (Supplementary Fig. 3). The model did not exhibit the post-inhibitory rebound excitation observed in the rodent recordings. L2/3 PCs also exhibited dual-peak excitation that preceded L5 PC responses, with peaks occurring around 10-15ms and 45ms, and excitation amplitude showing dose-dependent effects (Supplementary Fig. 4).

**Figure 3.**
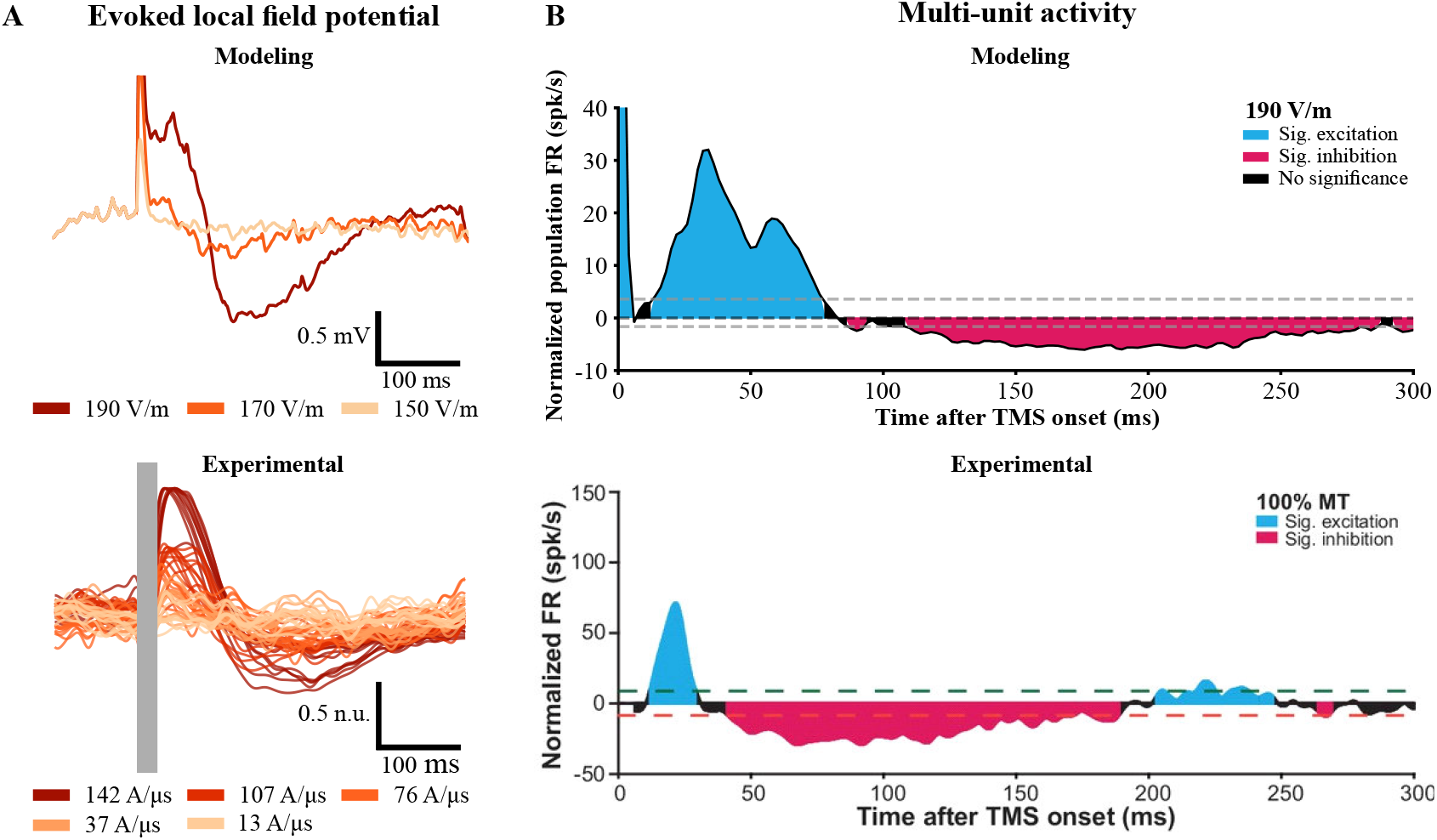
Comparison of model and experimental TMS-induced evoked response. **(A)** TMS evoked local field potential (LFP). Top: simulated evoked LFP signals recorded at L5 of the cortical column model. Bottom: evoked potential recorded in NHP, adapted from [37]. 25ms after the pulse onset was shaded in gray due to the strong pulse artifact from the experiment. **(B)** Evoked multi-unit activity of L5 neurons. Top: normalized average population FR of L5 PC (2ms time bins). Bottom: population average of normalized multiunit FR in layer V of rodent caudal forelimb area (CFA) in response to monophasic single pulse TMS at 100% motor threshold (MT), adapted from [38]. Dashed lines, significance thresholds determined by the 2.5 or 97.5 percentile of the baseline normalized FR [38].

**Figure 4.**
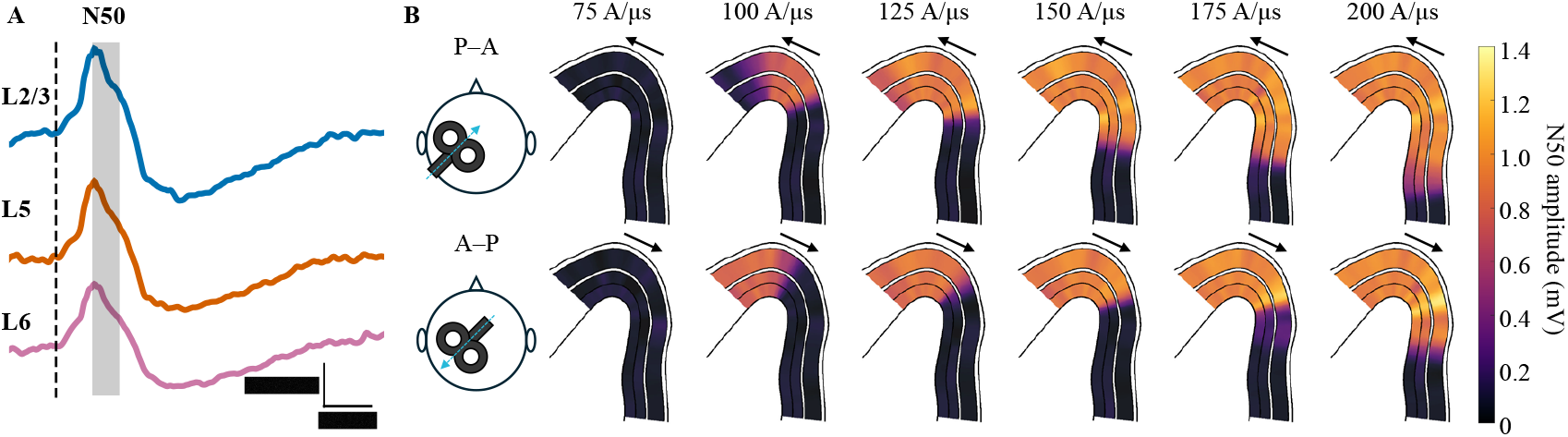
Spatial distribution of N50 varies with TMS intensity, shown in a cross-section of the hand knob. **(A)** Example LFPs computed at L2/3, 5 and 6 of a cortical column embedded in the anterior lip of the precentral gyrus in response to P-A TMS at 150 A/μs. N50 amplitudes are extracted in the time window of 40-70ms (shaded) from TMS onset (dashed line). **(B)** Spatial distribution of N50 amplitude on analysis planes through pre-central gyrus at different stimulation intensities. Arrows indicate direction of the initial phase of TMS induced electric field waveform.

Next, we systematically scaled the conductance of GABAB receptors at inhibitory synapses from SST to excitatory neurons by 0.5, 1.3 and 2.0 to examine effects on evoked responses. Increasing GABAB receptor conductance suppressed L5 PC FR during the 50–100 ms after stimulation (Supplementary Fig. 5), causing earlier onset and longer duration of the inhibitory phase.

We simulated the spatial pattern of TMS evoked responses by embedding the cortical column in an E-field head model at 30 locations arranged along a single cross-section of the left motor hand knob in M1. The TMS coil was positioned over the hand knob and oriented 45 degrees relative to the midline, corresponding to the lowest threshold configuration for evoking motor responses. Monophasic pulses were applied in two current directions (P–A and A–P) at six intensities ranging from 75 to 200 A/μs in 25 A/μs increments. We quantified the percentage of PCs directly activated by TMS in layers L2/3, L5, and L6 from each cortical column. Previous modeling work demonstrated that the activation pattern of isolated morphologically-realistic neuron models during monophasic stimulation is layer-specific and depends on the current direction [10] (Supplementary Fig. 6 right panel). For the same current direction, the lowest thresholds occurred at the gyral crown and lips. Our cortical column model replicated the layer-specific and spatial distribution of direct PC activation observed in previous studies, with higher percentages of direct activation at the gyral crown and lips as stimulation intensity increased (Supplementary Fig. 6 A and B). Switching from P–A to A–P monophasic TMS produced a pronounced anterior shift in direct activation percentages for L2/3 and L6 neurons below 125 A/μs, while this shift in L5 was only evident at 75 A/µs. These directional differences diminished at higher intensities (>125 A/μs) as activation approached saturation levels across all layers (Supplementary Fig. 6 C). Direct activation of model PCs occurred across the 30 cortical columns at the lowest tested intensity of 75 A/μs (Supplementary Fig. 7 and Table S2), representing the minimum activation threshold in our model.

To investigate the spatial distribution of TMS-evoked responses, we analyzed the early component of evoked LFPs (N50) recorded from the 30 cortical columns at three electrode locations (L2/3, L5, and L6) (Fig 4. A). The simulations revealed dose-dependent and spatially distinct N50 component patterns in response to P–A and A–P stimulation (Figure 4B). At low intensity (75 A/μs), N50 components were absent across the precentral gyrus in either direction. P-A stimulation elicited N50 responses that emerged first at the crown and posterior lip, subsequently expanding along the crown and posterior wall with increasing intensity. A-P stimulation produced N50 components primarily at the anterior lip, with the spatial distribution shifted anteriorly compared to P-A stimulation.

## Discussion

We developed a computational cortical column model to investigate the mechanisms underlying temporal responses to TMS. Our model demonstrated that single-pulse TMS evoked a characteristic response consisting of short-latency excitatory activation followed by prolonged inhibition, with both the strength of excitation and duration of inhibition scaling with stimulation intensity. Embedding this cortical column within a realistic head model revealed the spatial distribution of TMS-evoked responses and its dependence on the TMS induced current direction, providing mechanistic insights into the neural dynamics underlying cortical stimulation.

### Model Validation

Our cortical column model successfully reproduced key features of experimentally observed TMS responses, providing strong validation of the computational framework. The model-generated local field potentials exhibited dose-dependent N50 components that closely matched TEP recorded from NHPs [47], demonstrating the model’s capability to capture the essential electrophysiological signatures of cortical responses to TMS. Furthermore, multi-unit activity from L5 PCs exhibited multiphasic patterns consistent with experimental recordings from rodent motor cortex [48] and other brain regions of NHPs and cats [49– 51], including the characteristic early excitation followed by prolonged suppression. However, the model did not reproduce the post-inhibitory rebound excitation observed in vivo. This discrepancy likely stemmed from our model’s limited implementation of thalamic circuitry and lack of cortico-thalamic feedback mechanisms, which are crucial for generating rebound excitation elicited by cortical stimulation [52–54]. These findings demonstrated how multilayered cortical circuitry transforms the initial TMS-induced excitation into the characteristic sequence of early activation followed by prolonged inhibition seen in experimental recordings.

### Alternating Direct and Indirect Activation Drives Excitatory Response

Our model highlighted distinct temporal dynamics in directly versus indirectly activated cortical neurons. At 190 V/m stimulation intensity, we observed layer-specific patterns of direct activation, with L5 pyramidal cells showing the highest proportion of direct activation (63.85%), followed by L2/3 pyramidal cells (27.5%), while L6 pyramidal cells were predominantly activated indirectly (97.95% indirect activation).

This differential activation pattern illustrated that TMS effects unfold through a cascade mechanism where directly stimulated neurons trigger secondary activation of indirectly stimulated populations through synaptic connections. The temporal dynamics of these two populations demonstrate how relatively sparse direct activation can generate robust population-level responses that engage the majority of cortical neurons. Specifically, the network propagation effects amplify the initial direct response, with indirectly activated neurons contributing to sustained population activity during the early excitatory phase.

### Temporal Structure of Early Excitatory Response

Our model produced a novel prediction regarding the temporal structure of the early excitatory MUA: rather than a single peak of activation, we observed dual-peak dynamics across multiple cortical layers. L5 PCs exhibited two distinct peaks occurring at approximately 30ms and 60ms post-stimulation, while L2/3 PCs showed an earlier temporal sequence with peaks at ∼10-15ms and ∼45ms. Analysis of directly and indirectly activated neuronal populations demonstrated that these peaks correspond to temporally segregated contributions from different activation mechanisms.

Notably, the L2/3 PC peaks were predominantly driven by neurons that were not directly activated by TMS, indicating that indirect activation through synaptic transmission generated temporally structured responses that preceded the L5 PC responses. TMS-induced activity propagates through layer-specific synaptic networks with characteristic delays. The temporal sequence from L2/3 to L5 responses revealed how cortical circuits transform the initial TMS stimulus into layer-specific patterns of direct and indirect activation.

These computational predictions showed intriguing correspondence with human TMS-EEG studies that consistently reported early TEP components including N15 (∼15ms), P30 (∼30ms) and later positive components P50-P60 (∼50-60ms) following TMS at M1 [55,56]. The L2/3 PC early peak at 10-15ms aligned closely with the experimentally observed N15 component, while the L5 PC early peak at 30ms corresponded to the P30 component. The later peaks across both layers (45ms in L2/3, 60ms in L5) corresponded to the P50-P60 components. This temporal correspondence can be mechanistically plausible given that both EEG and LFP signals arise from the same fundamental process—mass synaptic inputs to geometrically aligned PCs—suggesting that our model may reflect the underlying synaptic activity patterns that generate the experimentally observed TEP components [57–61].

This dual-peak multi-unit activity differed from the single peak typically reported in experimental studies such as [48], which recorded from 7 neurons. The discrepancy likely reflects insufficient sampling in experimental recordings to resolve the temporal dynamics of population responses. Our comprehensive modeling approach, capturing activity from all 2,180 L5 PCs, enables detection of subtle temporal features. Additionally, extracellular and single-unit recordings may exhibit sampling bias toward larger neurons that are more sensitive to E-fields and thus more likely to be directly activated, potentially obscuring the contribution of indirectly activated populations to the overall response.

### Spatial Distribution and Coil Orientation Effects

Our multiscale model incorporating 30 cortical columns across precentral gyrus demonstrated spatially distinct patterns of TMS-evoked responses that depended on both E-field magnitude and current direction. The early components of TMS responses (N50) occurred preferentially at the lips and crown of the motor hand knob, where E-field intensity was highest due to proximity to the TMS coil. Despite better alignment of cortical columns to the current direction at the gyral walls, these regions exhibited weaker responses due to substantially lower E-field intensities resulting from their greater distance from the coil.

These findings are consistent with previous modeling work by Aberra and colleagues [10], confirming that TMS effects reflect the combined influence of E-field magnitude and cortical column orientation. The directional sensitivity we observed, with P–A stimulation producing responses at the crown and posterior regions while A–P stimulation preferentially activated anterior areas.

The dose-dependent emergence and spatial expansion of the N50 component further demonstrated how stimulation intensity determines not only the magnitude but also the spatial extent of cortical activation, with implications for the specificity and spread of TMS effects in therapeutic applications.

### TMS thresholds

In human TMS experiments, motor thresholds, measured with EMG, typically correspond to peak cortical E-fields from 60 to over 100 V/m [62–64], while modeling studies using isolated neuron models have demonstrated even higher thresholds [10,12,65,66]. Our detailed cortical circuit model, which includes endogenous activity and synaptic connectivity, yielded lower thresholds compared to quiescent cells in previous modeling studies [10,12]. At 150 V/m, approximately 25% of L5 PCs were directly activated by single monophasic TMS pulse, whereas the same PC models under quiescent conditions had minimum thresholds exceeding 180 V/m [10,12]. At 190 V/m, the cortical circuit model produced a robust early TEP component (N50). While these values may appear high compared to experimentally identified values, empirical evidence demonstrates significant local electric field variations at the microscale level. A recent tACS study in NHP has shown layer-specific conductivity variations within the gray matter of V1, with layers 2/3 exhibiting reduced electrical conductivity that resulted in higher local E-fields [67]. This finding suggested that microscale tissue heterogeneity can create substantial local field amplification compared to macroscopic calculations. Additionally, microstructure modeling studies have suggested potential for electric field modifications at the cellular level due to tissue heterogeneity [68,69], though the magnitude of these effects remains debated [70,71]. These findings indicated that the effective E-field experienced by individual neurons can differ substantially from macroscopic averages, supporting the physiological relevance of the field strengths used in our model.

### Limitations and Future Directions

Several limitations of the current model should be acknowledged. First, our implementation focused on cell types and morphologies that are most sensitive to TMS-induced E-fields, potentially overestimating the direct activation percentage compared to the full diversity of cortical neurons. Future iterations could incorporate a broader range of neuronal subtypes with varying sensitivities to electric stimulation.

Second, the model included only feedforward thalamic inputs and lacks the cortico-thalamo-cortical feedback loops that are known to contribute to cortical rebound response [52–54]. This limitation likely explains the absence of post-inhibitory rebound excitation observed in some experimental recordings and represents an important target for future model enhancement.

Third, our LFP calculations did not account for volume-conducted signals from distant sources, and the current implementation lacks cortical-to-cortical connections between columns within the head model. Incorporating these features would provide a more comprehensive representation of the distributed cortical responses to TMS and enable investigation of inter-regional connectivity effects. A key advancement would be incorporating volume conduction forward models to predict scalp EEG recordings in humans following TMS. By coupling our cortical column responses with realistic head models that account for cerebrospinal fluid, skull, and scalp conductivity, we could directly simulate the TEP recorded at the scalp level [57–61]. This would enable quantitative comparison of our dual-peak FR predictions with the temporal structure of human TEP components, potentially resolving the mechanistic origins of N15, P30, and P50-P60 components observed experimentally.

Finally, while used to simulate human TMS, our model incorporates neuronal morphologies reconstructed primarily from animal studies and validation data from rodent and NHP experiments. Direct experimental validation of these predictions in humans remains challenging due to ethical and technical constraints that prevent comparable detailed LFP and single-unit recordings during TMS. This cross-species modeling approach is justified by the fundamental conservation of cortical circuit organization and synaptic connectivity patterns across mammalian species [72,73]. As human intracranial recording data from TMS studies become increasingly available through clinical research opportunities, these datasets will provide crucial validation for refining human-specific cortical circuit models and enhancing the clinical translation of computational TMS predictions.

Despite these limitations, our model provided mechanistic insights into TMS-evoked cortical dynamics and establishes a validated computational framework for investigating the neural circuit basis of non-invasive brain stimulation effects. Future developments incorporating more comprehensive thalamic connectivity, inter-cortical connections, and volume conduction effects will further enhance the model’s utility for understanding and optimizing TMS protocols in both research and clinical applications.

## Acknowledgments

This work was supported by National Insitute of Biomedical Imaging and Bioengineering (NIBIB) and Collaborative Research in Computational Neuroscience (CRCNS): R01EB034143. We also acknowledge Research Computing and Minnesota Supercomputing Institute (MSI) for providing computational resources.

