## Supplemental Material for "Simulation of evoked responses to transcranial magnetic stimulation using a multiscale cortical circuit model"

### Supplementary Materials

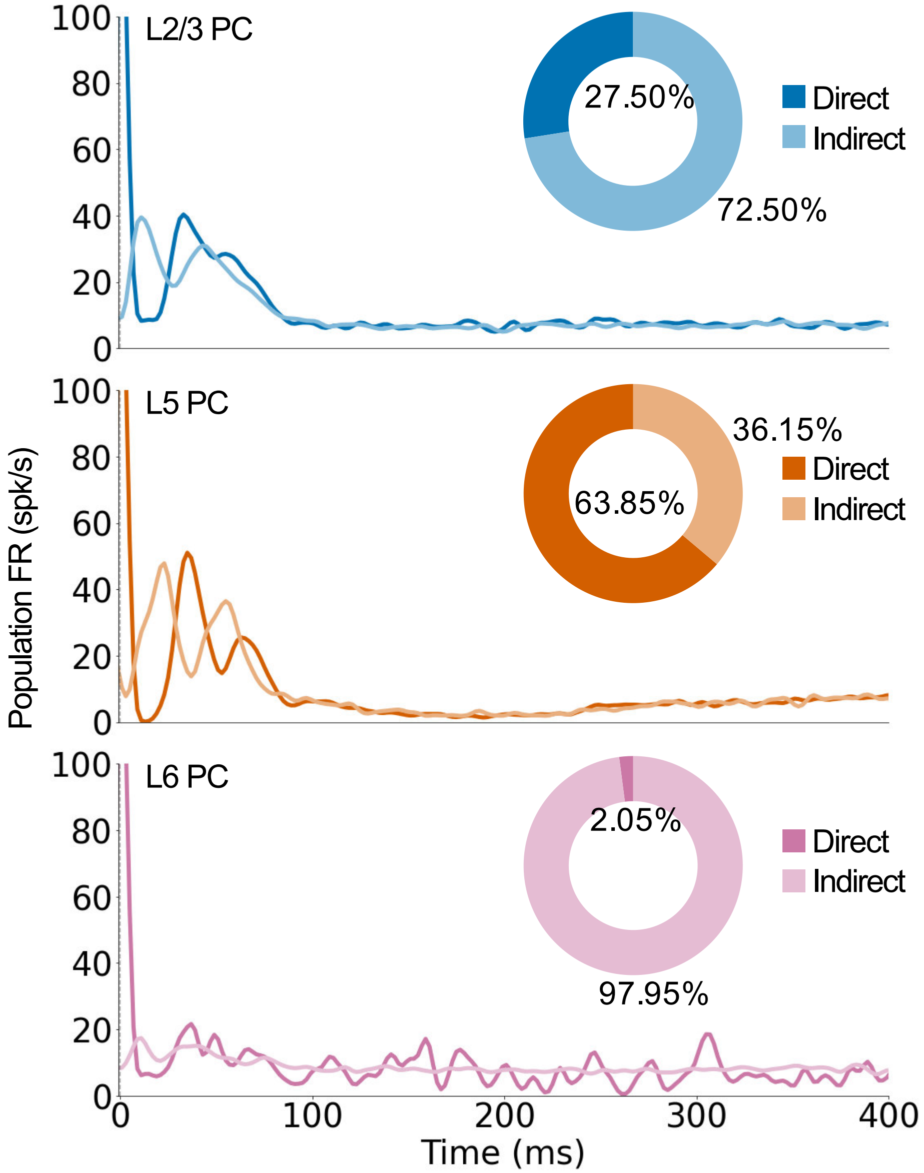


**Figure 1. Evoked response of PC in the cortical column at 190 V/m.** Population FR of PC in each layer. Cells in each population were categorized into directly activated (direct) and indirectly activated group (indirect) by the single pulse TMS at 0 ms. Pie charts show the proportion of direct and indirect groups in each population.

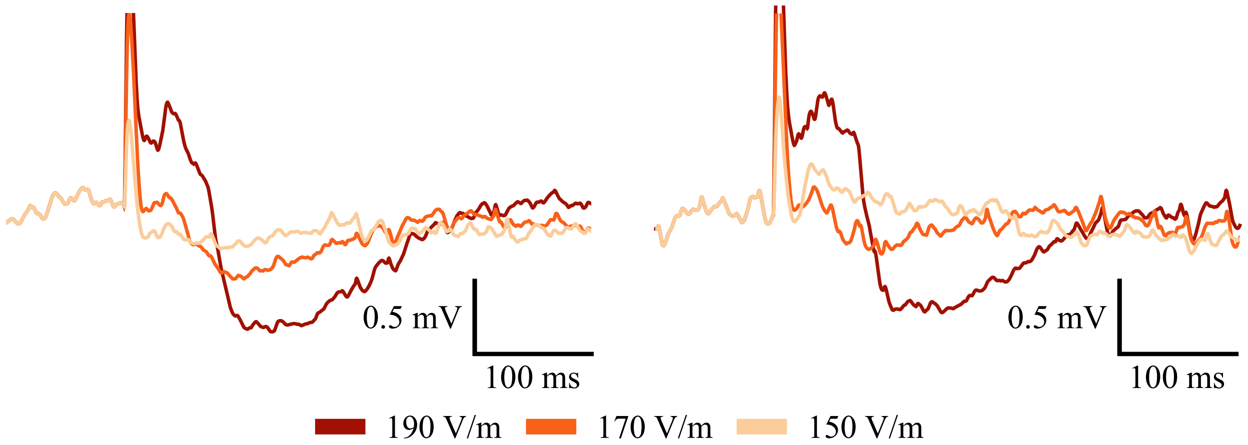


**Figure 2. Two additional independent trials of simulated TMS-induced evoked response with.** TMS evoked local field potential (LFP) recorded at L5 of the cortical column model with two sets of random seeds.

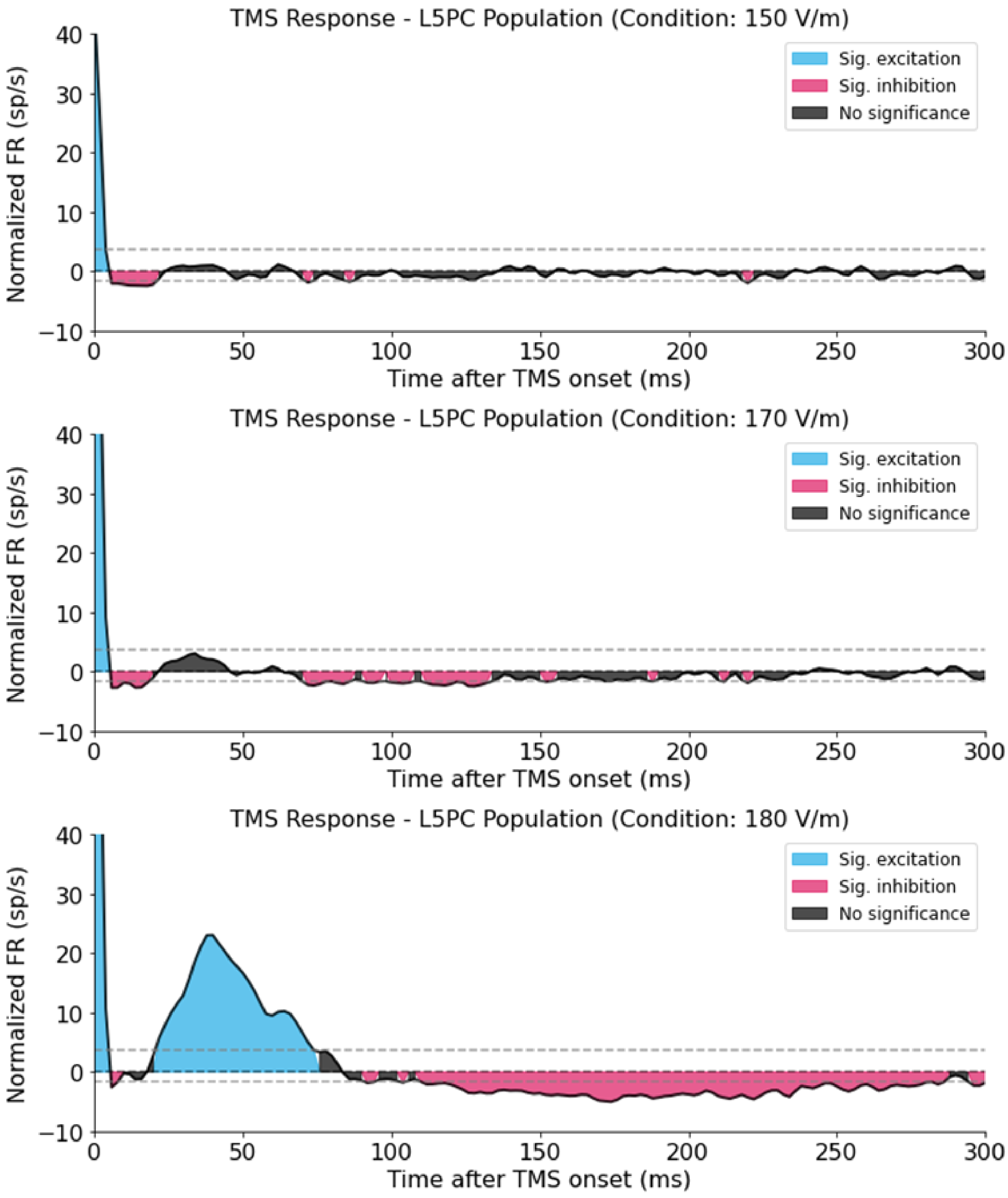


**Figure 3. Evoked multi-unit activity of model L5 PCs at different stimulation intensities.** Normalized average population FR of L5 PC (5ms time bins) at 150, 170 and 180 V/m, same approach as in main Fig. 3.

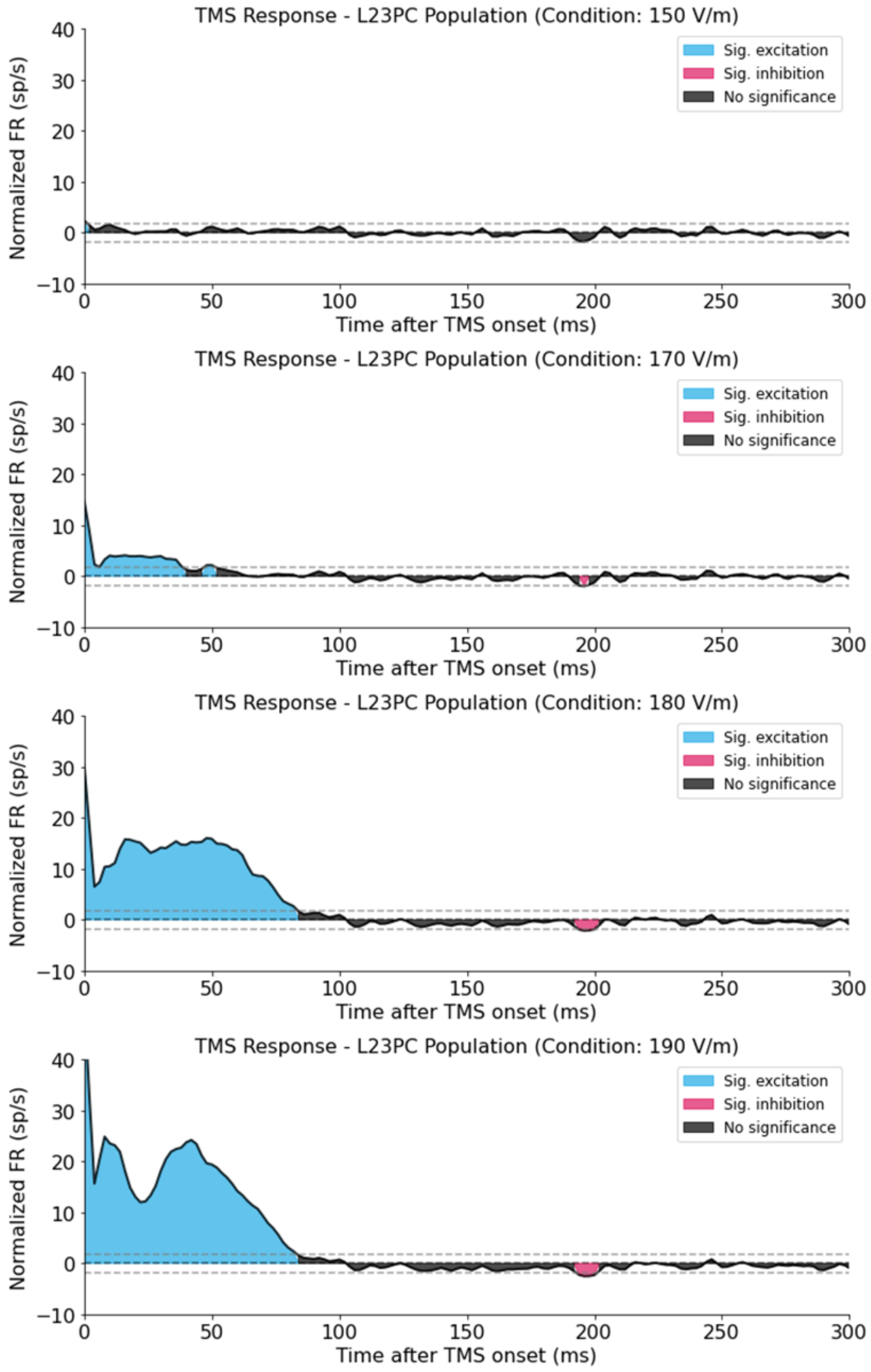


**Figure 4. Evoked multi-unit activity of model L2/3 PCs at different stimulation intensities.** Normalized average population FR of L2/3 PC (5ms time bins) at 150, 170, 180 and 190 V/m, same approach as in main Fig. 3.

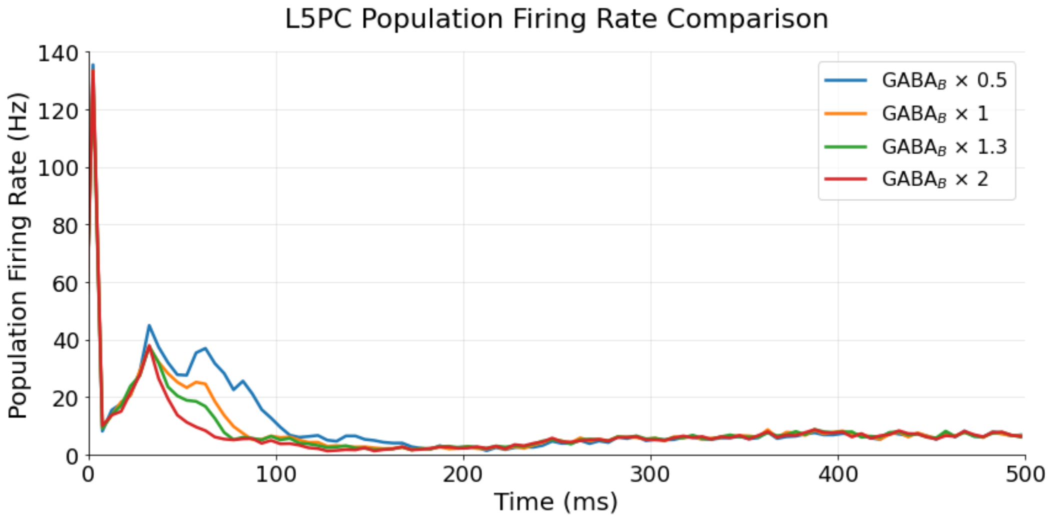


**Figure 5. Model L5 PC responses to TMS-induced uniform E-field for different levels of GABA$_B$ synaptic strength.** PSTH response at 200V/m and the proportion of GABA$_B$ conductance is scaled by 0.5, 1, 1.3 and 2. The population FR was averaged across all L5 PCs. The bin width is 5ms and the TMS onset is normalized to 0ms.

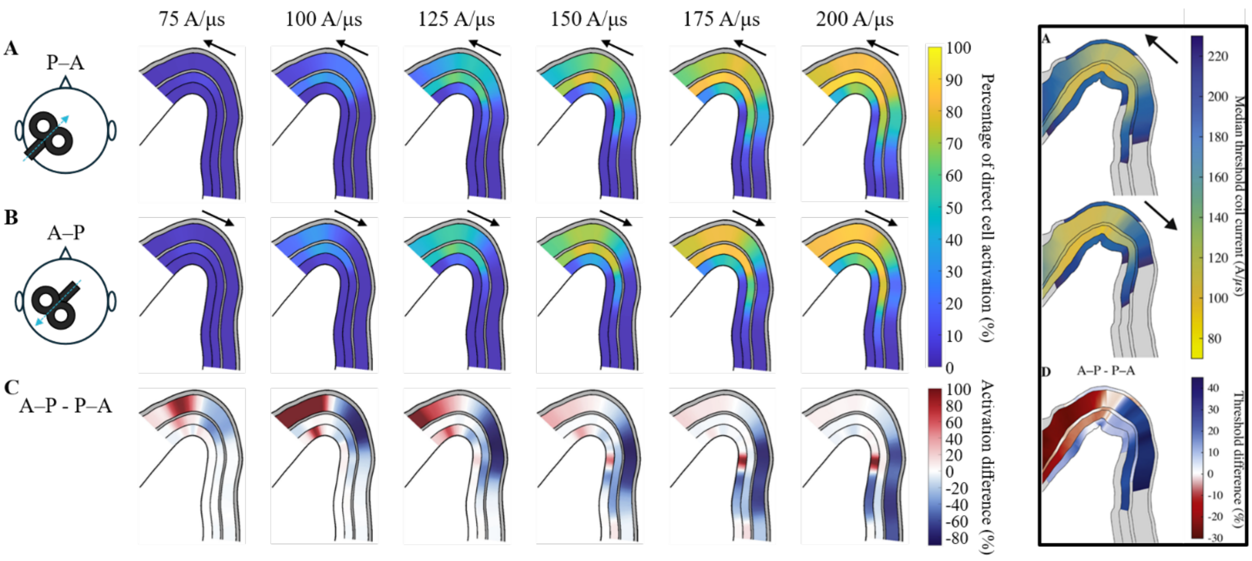


**Figure 6. Layer-specific spatial distribution of direct activation of PCs varies with TMS direction, shown in a cross-section of the hand knob.** The percentage of PCs directly activated by monophasic TMS in L2/3, 5 and 6 for (A) P-A and (B) A-P coil orientations at different stimulation intensities. (C) Percent difference in direct activation of PCs between P-A and A-P current directions. L1 and L4 are colored gray because no neurons are present in these two layers. The right panel shows median thresholds for L1-L6 on the plane through the precentral gyrus for monophasic stimulation in the P-A and A-P directions, adapted from [1]. PCs in L2/3, L5, and L6 from [1] were isolated, and their morphologies are identical to those used in the cortical column model shown in the left panel.

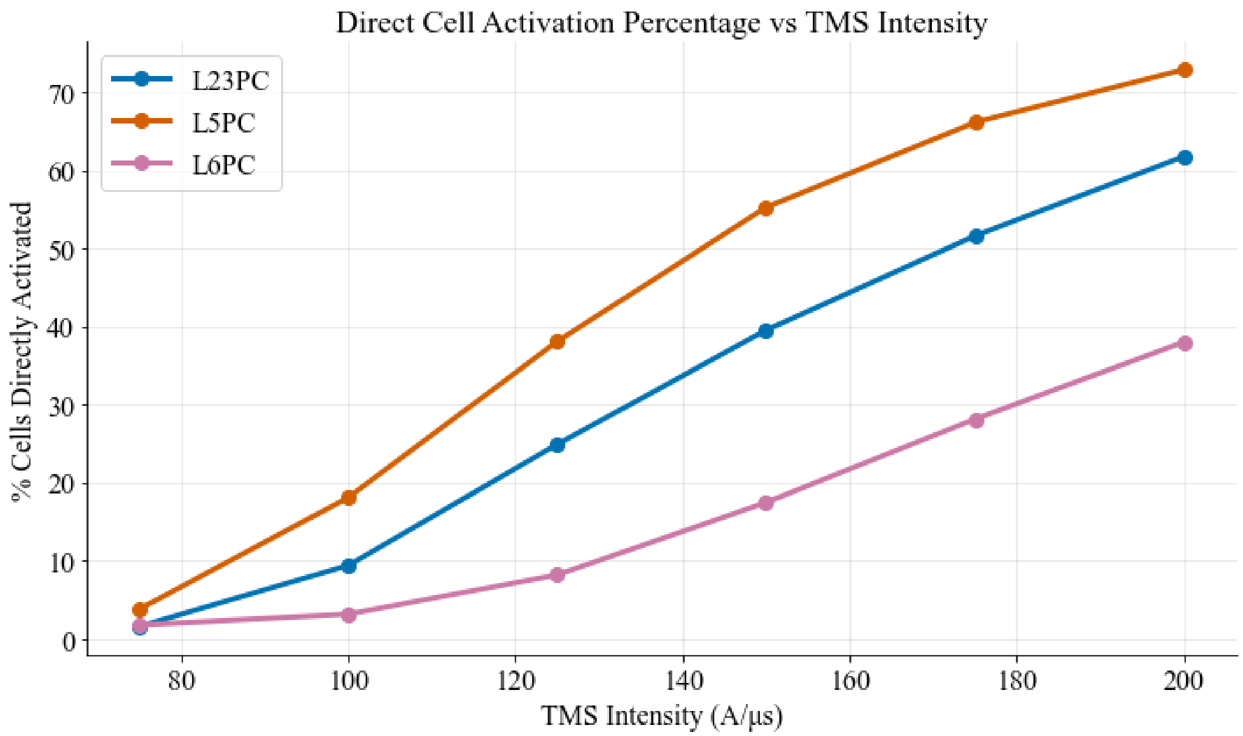


**Figure 7. Direct PC activation percentage of 30 embedded cortical columns at different TMS intensities under P-A TMS.** The percentage of direct PC activation was calculated based on the total number of cells across all 30 cortical columns (L2/3 PC: 104,400; L5 PC: 65,400; L6 PC: 70,200). The exact values at each data point are shown in Table S2.

**Table S1. Direct PC activation percentage of a cortical column (Uniform E-field)**

| | Direct Cell Activation Percentage (%) | | |
|---|---|---|---|
| TMS Intensity (V/m) | L2/3 PC | L5 PC | L6 PC |
| 150 | 2.41 | 25.50 | 1.84 |
| 170 | 8.65 | 51.80 | 1.84 |
| 180 | 16.50 | 59.90 | 1.92 |
| 190 | 27.50 | 63.85 | 2.05 |

**Table S2. Direct PC activation percentage of embedded 30 columns (P-A TMS)**

| | Direct Cell Activation Percentage (%) | | |
|---|---|---|---|
| TMS Intensity (A/μs) | L2/3 PC | L5 PC | L6 PC |
| 75 | 1.54 | 3.75 | 1.75 |
| 100 | 9.36 | 18.06 | 3.12 |
| 125 | 24.87 | 38.04 | 8.14 |
| 150 | 39.49 | 55.22 | 17.44 |
| 175 | 51.57 | 66.11 | 28.11 |
| 200 | 61.75 | 72.85 | 37.98 |
